# Characterization of calcifications in posterior horn of human meniscus using micro-computed tomography

**DOI:** 10.1101/2025.11.07.686933

**Authors:** Ville-Pauli Karjalainen, Iida Hellberg, Aleksandra Turkiewicz, Bijay Shakya, Nodira Khoshimova, Eeva Nevanranta, Shuvashis Das Gupta, Khaled Elkhouly, Amanda Sjögren, Mikko A.J. Finnilä, Patrik Önnerfjord, Velocity Hughes, Jon Tjörnstrand, Martin Englund, Simo Saarakkala

## Abstract

**Objective:** Meniscal calcifications are associated with meniscal degeneration and osteoarthritis (OA). We investigated micro-computed tomography (µCT) imaging for identification of calcification patterns caused by basic calcium phosphate (BCP) and calcium pyrophosphate (CPP).

**Design:** Posterior horns of human medial and lateral menisci from 19 individuals with medial compartment knee OA and 21 deceased donors were imaged with high-resolution µCT. Raman spectroscopy characterized the calcification types from histological sections, adjacent to µCT piece. Qualitative and quantitative analysis, visualization, and grading were performed using 3D µCT images of meniscal calcifications.

**Results:** Different calcification patterns were observed with BCP and CPP. BCP was found at the borders of meniscal tissue and inside complex tears or fibrillation. In contrast, CPP accumulated as circumferential rod-like structures between collagen bundles, mainly inside the meniscal tissue, and lacked the porous 3D structure observed in BCP calcifications. Quantitatively, CPP samples had a higher calcification volume compared to BCP with a geometric mean ratio of 14 (95%CI; 3,73), and larger particle sizes with a ratio of 7 (95%CI; 2,8). Additionally, BCP calcifications had an organized porous structure with a closed porosity range of 6–19, while a similar structure was not seen in CPPs (range 1–3.5).

**Conclusions:** We qualitatively and quantitatively identified volumetric and morphological differences in the calcification deposition patterns between BCP and CPP calcifications in human meniscus. The calculated differences may help distinguish the calcification types with *in vivo* imaging modalities in the future, as well as provide a better understanding of their role in OA.

## 1. Introduction

Calcifications in the meniscus are suggested to be an essential part of meniscal degeneration and osteoarthritis (OA)^1–3^. Meniscal calcifications propose a predisposing factor for cartilage lesions and OA progression, and therefore, this ectopic mineralization should be taken into consideration when developing OA treatments^4,5^. Currently, there are no effective disease-modifying drugs for OA, but some drugs have shown potential in inhibiting the pathological calcification^6–8^. Understanding the different calcification deposition patterns caused by different calcification types may increase the knowledge regarding calcification processes in OA and other diseases with crystal deposition involvement, which could, consequently, help in future drug development.

Calcium pyrophosphate (CPP) and basic calcium phosphate (BCP), such as hydroxyapatite are the most common types of meniscal and articular cartilage calcifications. BCP calcifications have been associated with OA and inflammation, while CPP is related to chondrocalcinosis and aging^8–11^. Methods such as Raman spectroscopy and scanning electron microscopy with energy dispersive analysis (SEM-ED) can differentiate BCP and CPP calcifications, but they can only analyze small samples. Furthermore, imaging with multi-energy computed tomography (MECT) combined with photon-counting detectors (PCD) has previously enabled the differentiation of calcifications in 3D with varying accuracy^8,12,13^.

We have previously shown that micro-computed tomography (µCT) can visualize and quantitatively analyze meniscal soft tissues and calcifications in 3D^5,14–17^. High-resolution µCT imaging enables non-destructive assessment of relatively large samples, provides accurate 3D information of the entire structure, and detects minute calcified particles that are often missed by conventional 2D techniques. We therefore propose that combining high-resolution µCT with Raman spectroscopy will allow identification of different calcification patterns with different calcification types.

This study aimed to utilize high-resolution µCT to quantitatively characterize and study the calcifications in human menisci *ex vivo*. The objectives were to volumetrically characterize and compare the deposition patterns of BCP and CPP crystal types in the medial and lateral posterior horns of osteoarthritic knees from total knee replacement (TKR) patients and donors without clinical knee OA.

## 2. Method

### 2.1 Tissue samples

This study was approved by the regional ethical review board at Lund University (Dnr 2015/39 and Dnr 2016/865, Dnr 2019/00323). The samples used in this study were obtained from the knee tissue biobank MENIX, located at Skåne University Hospital in Lund, Sweden. This biobank contains meniscus samples from two distinct groups: individuals with knee OA who underwent a total knee replacement (TKR) at Trelleborg Hospital and deceased adult donors (samples obtained within 48 hours post-mortem) with no known history of knee OA. All samples in the biobank were preserved at −80 °C within two hours of extraction.

The sample set of this study consisted of 80 meniscus samples: we selected both the medial and lateral menisci from a single knee from 19 individuals with end-stage medial compartment knee OA and 21 deceased adult donors without a known diagnosis of knee OA or rheumatoid arthritis. In the TKR group, the surgeon’s Outerbridge classification of knee joint cartilage was used to assess the primary compartment affected by OA. Individuals who underwent TKR with a medial grade of IV and a lateral grade lower than IV were included in the study. Additionally, the samples required a surgeon’s sketch revealing at least a part of the posterior horn of the medial meniscus to be remaining, to be considered eligible for the study. Macroscopic intactness of the deceased donor menisci was also required for inclusion in the study. Given that all available TKR recipients were older than 50 years, we applied the same age threshold for deceased donors.

### 2.2 Sample preparation

After receiving the meniscus samples from the MENIX biobank, they were thawed in phosphate-buffered saline (PBS). Supplementary material figures S1 and S2 show images of thawed donor and TKR menisci before any sample processing. In macroscopic images, the largest CPP calcifications can be seen by eye, circulating inside the meniscus, aligning with circumferential collagen fiber bundles (Figure S1). No cases of visible calcifications in the BCP samples were seen. The menisci were divided into two parts with a scalpel. The posterior horn with some part of the body was used in this study. The posterior horns were then fixed in 4% saline-buffered formaldehyde until they were dissected into several 5–10 mm thick pieces. The first piece was subsequently immersed in formalin, water, absolute alcohol, and xylene, then infiltrated with molten paraffin. No decalcification process was performed on the samples to preserve the calcifications. The processed samples were then manually embedded into paraffin blocks. Vertical and horizontal 4-µm-thick sections stained with Safranin O – Fast green, and hematoxylin and eosin (H&E) were used in the histopathological scoring of meniscal degeneration. In addition, vertical Alizarin red stained sections were used for visual inspection of meniscal calcifications. Furthermore, 5-µm-thick vertical sections were positioned on highly polished stainless steel Raman windows for Raman measurements. Before the Raman measurements, the paraffin was chemically dewaxed from each sample. The Raman spectral spectroscopy measurements are explained in more detail in our previous study^9^. The samples were then split into either BCP or CPP group based on the Raman results.

### 2.3 µCT imaging

Adjacent to the histological sections, the fixed µCT samples were first dehydrated in ascending ethanol concentrations (30%-50%-70%-80%-90%-96%-100%), treated with HMDS for 4 hours, and air-dried in a fume hood overnight. The image acquisition was performed using a desktop µCT device (SkyScan 1272, Bruker microCT, Kontich, Belgium) with the following settings: tube voltage 60kV; tube current 166µA; no additional filtering; isotropic voxel size 2.0µm; number of projections 2400; averaging 2 frames/projection; random movement 25 pixels; and exposure time 3500ms. NRecon software (Version 2.2.2.0 Bruker microCT) was used for image reconstruction. Two image reconstructions were performed for each sample with optimized settings and windowing for soft tissue and calcifications, producing two image stacks as an output. During image reconstructions, beam-hardening and ring-artefact corrections were applied together with pixel masking for defect pixels. Six out of 80 samples were omitted from further analysis due to image acquisition failure with µCT, resulting in a total of 74 samples.

### 2.4 Grading of meniscal calcifications

The calcifications in the menisci were graded between 0-5 from no calcifications to widespread calcifications from the 3D µCT images according to Hellberg et al^5^. Three individuals (VPK, IH, NK) graded the samples individually and then attained a consensus grade that was used as the final grade.

### 2.5 Micro-computed tomography analysis of meniscal calcifications

The whole µCT pieces were used for the analyses. A sharpening filter followed by thresholding, and despeckling were applied using CTAn software (Version 1.20.2 Bruker microCT). We calculated the total calcification volume and meniscus tissue volume using the full 3D image stacks. In addition, for each sample, we performed individual object analysis, which describes the 3D morphology of each individual calcification particle within each sample. It gives the following parameters as an output: number of particles, individual calcification particle volume, surface to volume ratio, sphericity, and closed porosity volume. Surface to volume ratio describes the complexity or roughness of calcification surface with higher value indicating more complexity. High sphericity indicates a sphere-like morphology, while lower values indicate rod-like or other more complex shapes.

### 2.6 Statistical analysis

SPSS (IBM SPSS Statistics, v. 29.0, NY, USA) was used for statistical analysis. For all parameters, we did a group-wise comparison between BCP and CPP groups using linear mixed models. All parameters were transformed into logarithms before calculating arithmetic mean for each meniscus which was used as outcome in the statistical models. The results were back transformed and thus, the difference between groups are expressed as ratios of geometric means. Used parameters were set as dependent variable and we used a random effect to account for multiple menisci from the same knee (menisci nested within persons). For total calcification volume and total number of particles, we adjusted for the meniscus logarithm tissue volume to account for differing sizes of meniscus. For closed porosity, volume of closed porosity was set as dependent variable, and the logarithm of total calcification volume was used as a fixed variable to account for differing total volumes of calcification. We did not adjust for any other covariates due to low number of menisci with CPP calcifications. We checked the assumptions of the linear mixed models using plots of residuals, QQ plots for normality of residuals and random effects and residuals vs fixed fitted values plot to assess linearity and homoscedascticity. We found no evidence of violation of the assumptions.

## 3. Results

### 3.1 Descriptive statistics on the study participants

Individuals with OA and deceased donors had an average age of 71 years, with similar height of 170 cm and half of them being females (Table I). The study subjects had an average age of 70.6 years with average age of 170 cm with half of them being females. (Table I).

**Table I.**
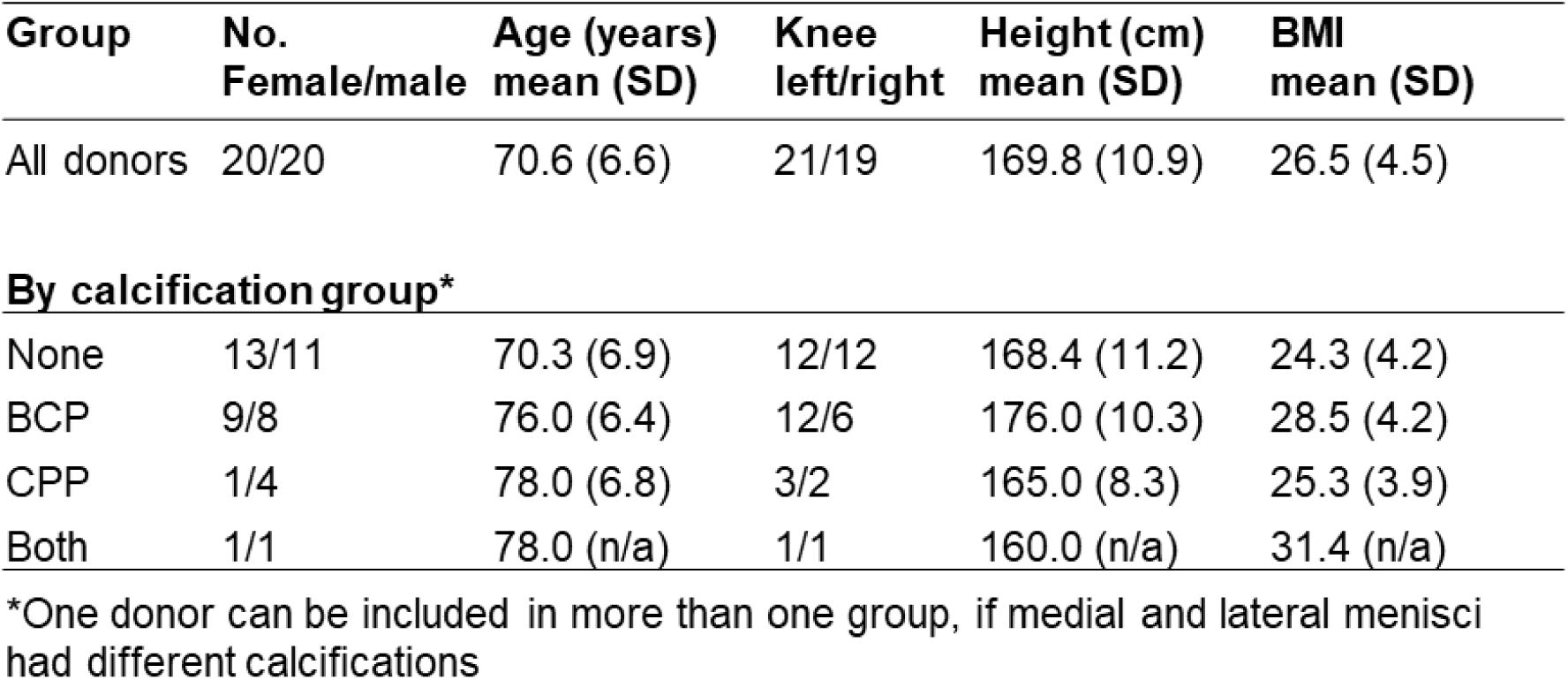
Descriptive statistics on all study participants and then divided into calcification types.

| Group | No.<br>Female/male | Age (years)<br>mean (SD) | Knee<br>left/right | Height (cm)<br>mean (SD) | BMI<br>mean (SD) |
| --- | --- | --- | --- | --- | --- |
| All donors | 20/20 | 70.6 (6.6) | 21/19 | 169.8 (10.9) | 26.5 (4.5) |
| **By calcification group\*** |  |  |  |  |  |
| None | 13/11 | 70.3 (6.9) | 12/12 | 168.4 (11.2) | 24.3 (4.2) |
| BCP | 9/8 | 76.0 (6.4) | 12/6 | 176.0 (10.3) | 28.5 (4.2) |
| CPP | 1/4 | 78.0 (6.8) | 3/2 | 165.0 (8.3) | 25.3 (3.9) |
| Both | 1/1 | 78.0 (n/a) | 1/1 | 160.0 (n/a) | 31.4 (n/a) |

\*One donor can be included in more than one group, if medial and lateral menisci had different calcifications

### 3.2 Qualitative assessment of meniscal calcification patterns from micro-computed tomography imaging and histology

Visual examination of the µCT images revealed different calcification patterns between BCP and CPP samples. In Figure 1, 3D µCT images show how BCPs are spread mostly on the surface, periphery and near the tears of meniscus. While few BCP particles seem to be inside the soft tissue in both 3D µCT and histology, they are in fact located inside complex tears or fibrillations of the meniscus. Similarly, Supplementary Video 1 shows BCP calcifications in an OA meniscus, mainly located on the surfaces and inside fibrillations. They appear mostly aggregated as punctate and small in size (5-100 µm in diameter), while larger clusters can be over 500 µm in the largest diameter and have characteristically sharp and pointy edges. Compared to the histology, not only larger volumes of calcifications are seen, but more diminutive calcifications can also be identified in 3D µCT.

**Figure 1.**
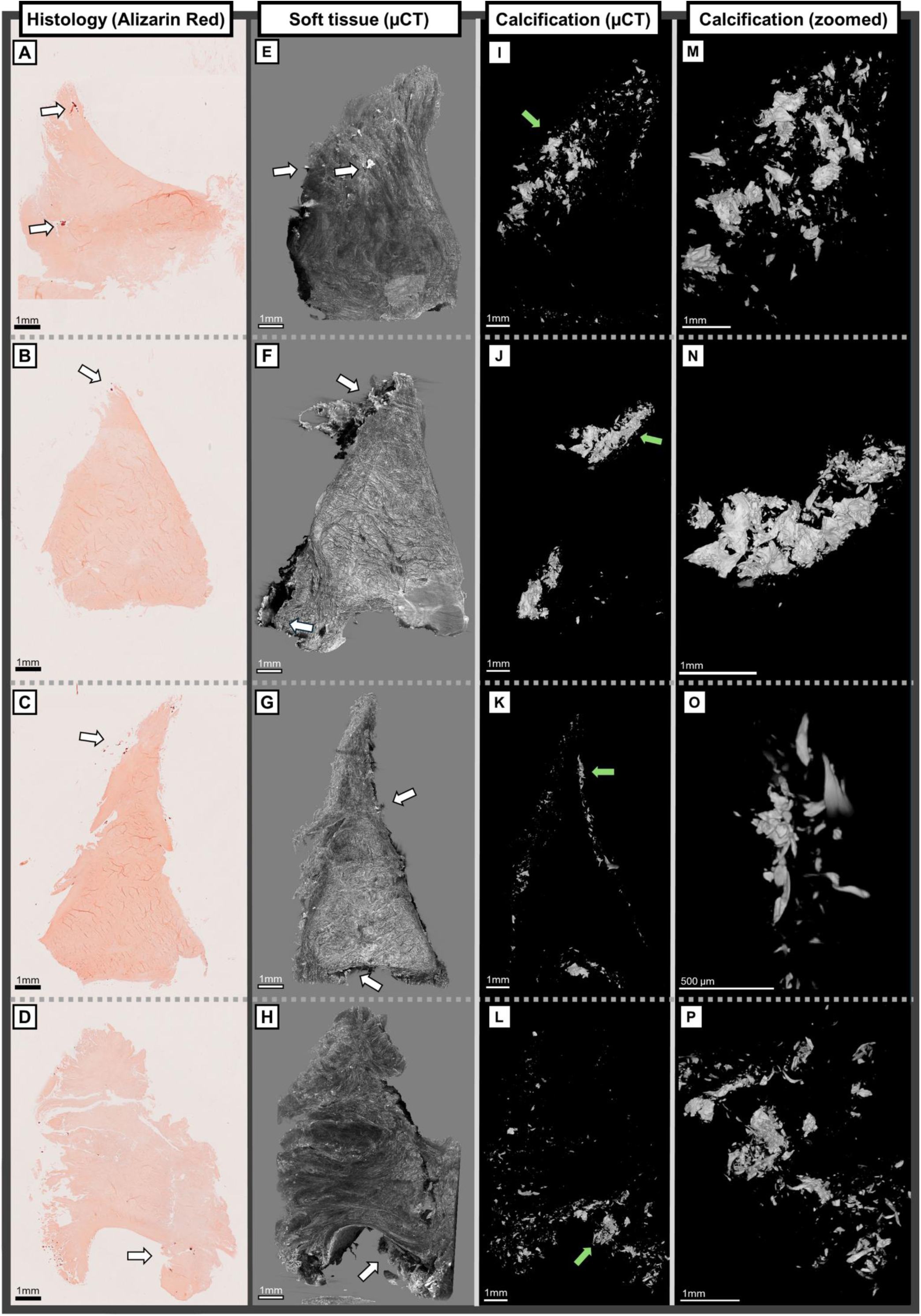
A, B, C, D) Overview of Alizarin red stained histological sections with basic calcium phosphate (BCP) calcifications from different meniscus samples. White arrows show areas with positive staining marking meniscal calcifications. E, F, G, H) 3D µCT images from meniscus soft tissue reconstruction show calcifications in white and meniscus tissue in gray. While some BCP particles seem to be inside the soft tissue in histology, from 3D µCT images it can be observed that they are located in the periphery of the meniscus, stuck inside complex 3D tears, or near fibrillations. Arrows highlight the calcifications in the image. I, J, K, L) 3D µCT images from meniscus calcification reconstruction. BCPs appear as punctate aggregates, measured 5-100µm in largest diameter, but can also accumulate into larger clusters of calcifications, sized even between 0.5 mm and 1 mm. The green arrows highlight the area of the zoomed image. (M, N, O, P) The zoomed calcification images show sharp morphology of the BCP calcifications, a likely contributor of tearing and degradation that advance the state of fibrillation while aiding the calcifications to bury in the surface fibrillations. [Suggested place for Supplementary Video 1]

CPP deposits appear mostly inside the meniscus tissue as long rod-shaped formations or ellipsoid-shaped clusters oriented along the circumferential collagen fibers (Figure 2). Supplementary videos 2&3 show meniscus samples with CPP calcifications covering most of the tissue. We observed that the cross-sectional diameter of these solid rods was commonly between 0.2 and 1 mm. The largest calcification clusters seemingly accumulate between the circumferential collagen fiber bundles, aligning their largest diameter with the fibers, while pushing and forcing the collagen to warp around them. The CPP rods can cover most of the meniscus from anterior horn to posterior horn, as seen in Supplementary Figure S1 where thick CPP calcification can be seen inside the meniscus through the surface macroscopically. Furthermore, few cases of long and hollow calcifications were observed (Supplementary Figure S3, Supplementary Video 3). Even smaller, ellipsoid-shaped calcifications align with the circumferential collagen orientation. However, when seen on the surface of meniscus, CPP calcifications are spread along the surface instead of along the circumferential collagen network. Additionally, CPP calcifications have generally smooth surfaces when appearing inside the tissue. Amorphous, less dense structures of CPP are seen near the cylinder-shaped calcifications and on the surface of the meniscus. Differences between solid rod-shaped and more amorphous CPP calcifications can be seen in Figure 2E, H, K.

**Figure 2.**
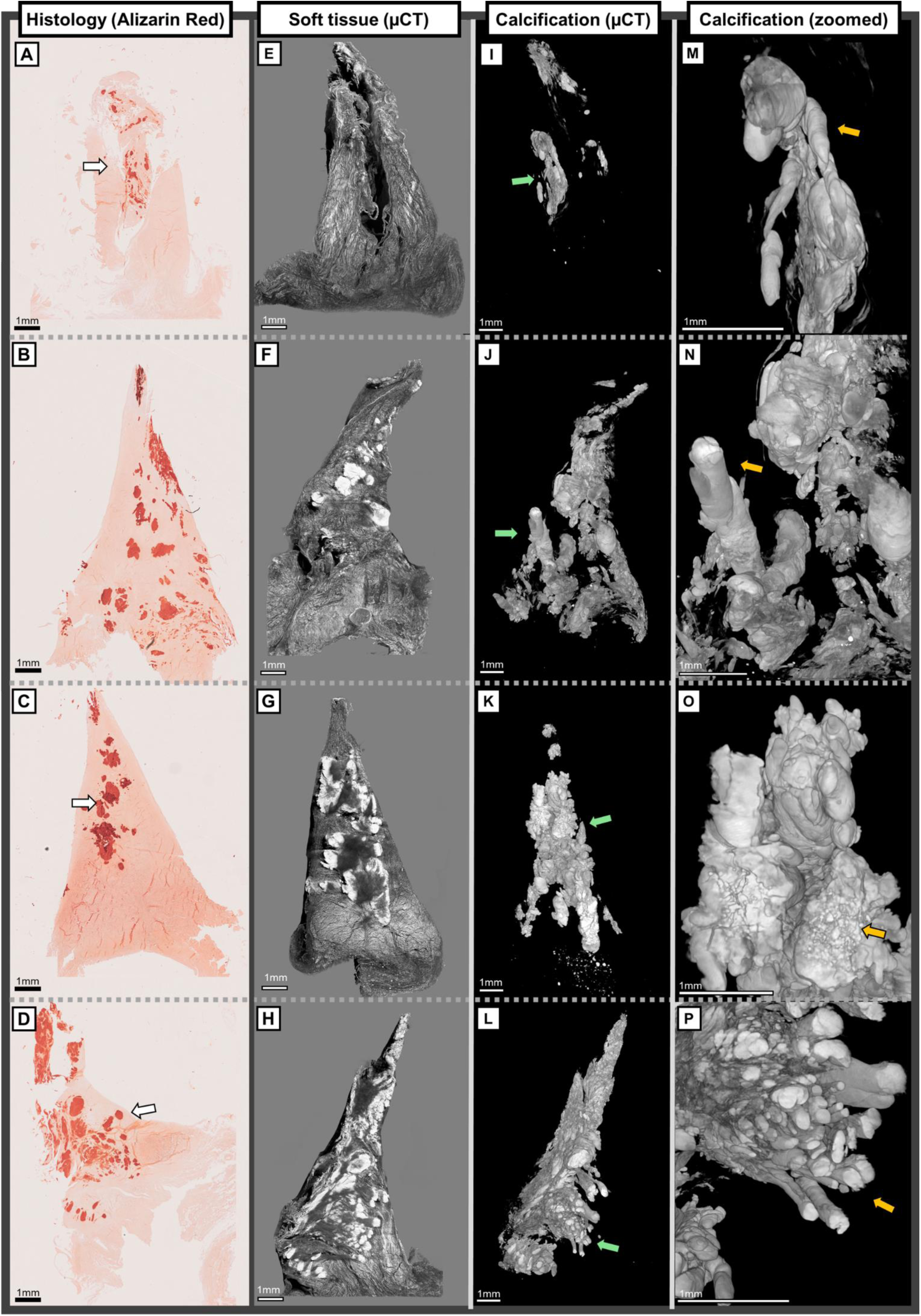
A, B, C, D) Overview of Alizarin red stained histological sections with calcium pyrophosphate (CPP) calcifications from four different samples. 2D sections show mostly spherical calcifications inside the tissue with more amorphous structures on the periphery and surface. E, F, G, H) 3D µCT images from meniscus soft tissue reconstructions show calcifications in white and meniscus tissue in gray. Similar to 2D histological sections, the calcifications cover large areas inside the meniscus. I, J, K, L) 3D µCT images from meniscus calcification reconstructions. As shown in 3D, long, rod-shaped calcification formations cover large areas inside the meniscus, while less dense, amorphous calcified structures are located on the surface and between the more solid calcifications. Green arrows show the zoomed area of calcifications. M, N) Smooth, rod-shaped calcifications align with the circumferential collagen fiber bundles inside the meniscus (orange arrow). O) Smooth aggregates of calcifications inside the tissue are accompanied by more amorphous calcification structures that are located on the tibial surface or between the solid calcifications (orange arrow). P) Multiple rod-shaped calcifications cover almost the whole meniscus (orange arrow) [Suggested place for Supplementary Video 2] [Suggested place for Supplementary Video 3]

Qualitative description of calcifications, cells, and proteoglycan content in the meniscus from histopathological sections are seen in Supplementary Material IV. Two example meniscus samples, one with both BCP and CPP calcifications and other with only BCP calcifications as identified with Raman together with Alizarin red, H&E, and Safranin O – Fast Green staining are seen in Supplementary Figures S4 and S5.

### 3.3 Porosity of calcifications

µCT inspection of the BCP calcifications revealed a porous, organized 3D structure (Figure 3A-D) that was consistently observed across all samples with BCP calcifications. While some pores were observed inside the CPP calcifications in µCT images, their amount was substantially lower than in BCP calcifications (Figure 3E-H). Supplementary Video 4 of a close-up BCP calcification with segmented pores shows how constant the pore structure is.

**Figure 3.**
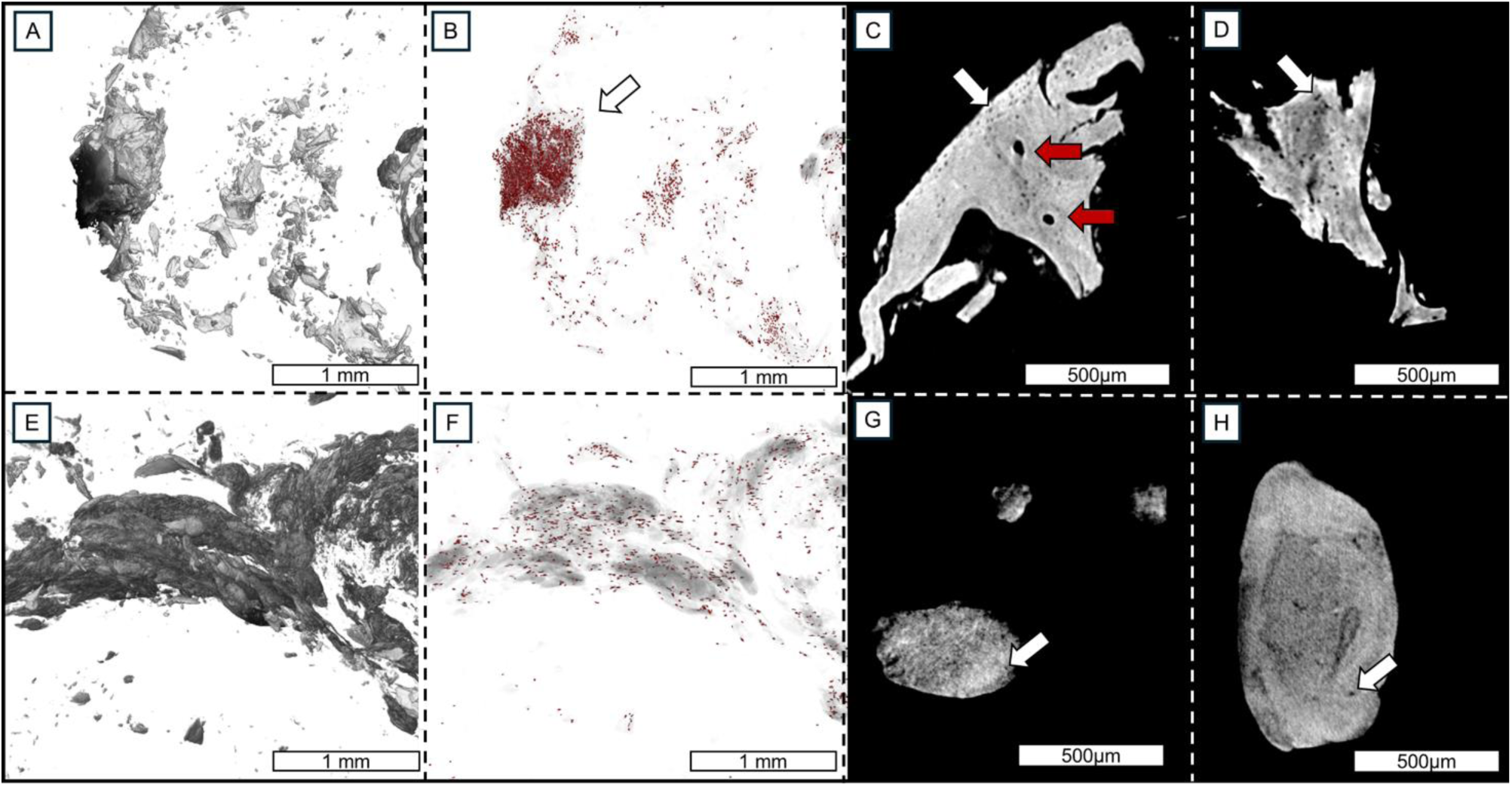
Closed porous structures inside the calcifications. A) 3D µCT image of a BCP cluster located in the outer periphery of meniscus. B) Lacunae-like pores inside the calcifications are depicted in red color. C) 2D cross-section of a BCP calcification cluster from Figure 3B, shows organized lacunae-like pores inside the BCP calcification (white arrows). Red arrows depict continuous, open channels– resembling Haversian canals in cortical bone–going through the calcification cluster. D) A similar 2D cross-section of a BCP cluster from a different sample shows the same porous and organized structure inside the BCP calcifications. E) CPP calcifications from the same sample as A). F) Closed porous structures appear in lower density than in BCP calcifications. G) 2D cross-section of a CPP calcification shows some porous structures inside the calcification, but to a lesser extent than in BCP calcifications (white arrow). H) 2D cross-section of a CPP cluster from a different sample shows a similar structure with only a few lacunae-like pores (white arrow). [Suggested place for Supplementary Video 4]

### 3.4 Grading of meniscal calcifications

The calcifications were graded in each meniscus sample from the 3D µCT images. As visualized in Figure 4, the median calcification grade was higher in individuals with OA compared to deceased donors [median (1^st^ quartile, 3^rd^ quartile)]: OA Lateral [3 (3,4)], OA Medial [4 (4,4)], Donor Lateral [1 (1,2.5)], Donor Medial [1 (0,2)]. Furthermore, even though donor menisci had lower median calcification grades compared to OA menisci, high grades were observed also in this group in samples with CPP calcifications. Eight donor samples had a grade 0, meaning that there were no visible calcifications in them.

**Figure 4.**
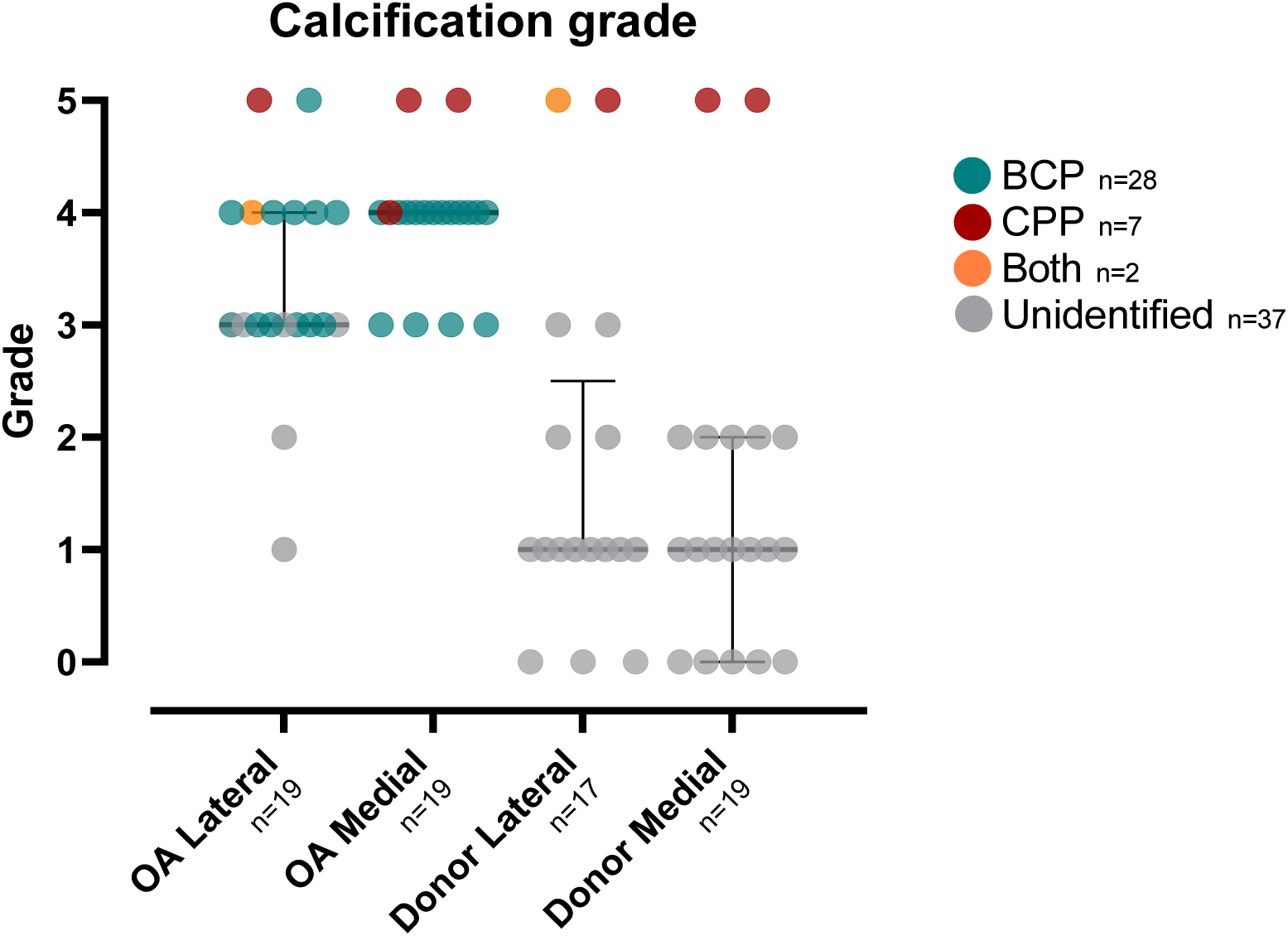
The association of OA status and joint compartment with calcification grade is shown with medians and their interquartile ranges. The calcification types are highlighted with different colors. The calcification grades were higher in individuals with OA compared to deceased donors. In addition, the menisci with CPP calcifications had higher calcification grades than samples with BCP calcifications. Only eight donor samples had a grade 0, meaning that there were no visible calcifications in them.

### 3.5 Quantitative 3D calcification analysis from micro-computed tomography images

BCP and CPP calcifications were quantitatively analyzed in 3D from the µCT images (Figure 5, Table II). The total calcification volume was higher in CPP compared to BCP (Figure 5A). Similarly, the total number of particles was greater in CPP than in BCP (Figure 5B). The average particle volume was also larger in CPP (Figure 5C). In contrast, the surface-to-volume ratio was higher in CPP (Figure 5D), while sphericity was greater in BCP (Figure 5E). Lastly, porosity was higher in BCP compared to CPP (Figure 5F). The total number of calcifications analyzed per meniscus is seen in Supplementary Figure 6.

**Figure 5.**
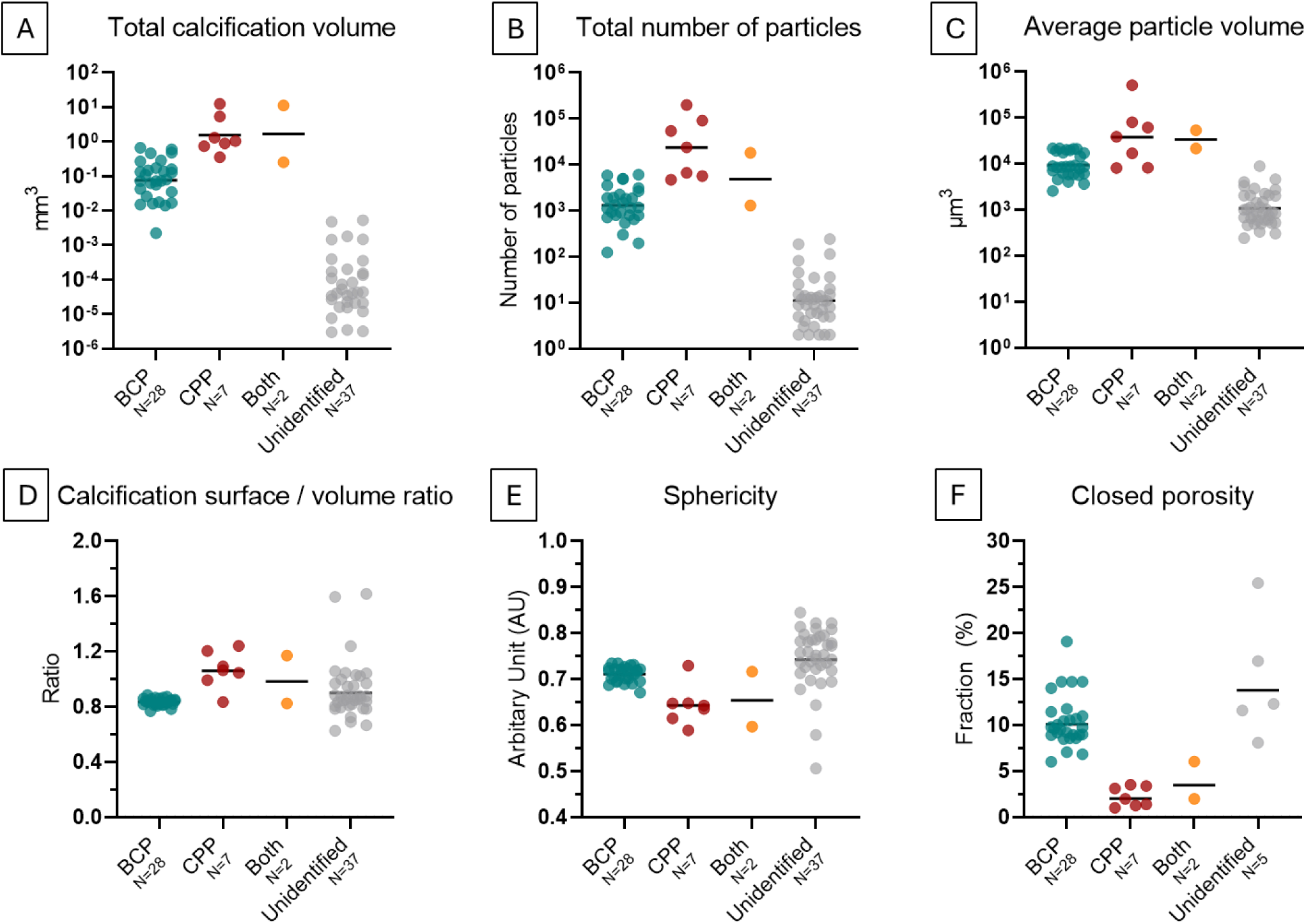
Quantitative 3D analysis of calcifications A) Geometric means of total calcification volume in each sample. B) Geometric means of total number of calcification particles. C) Geometric means of average particle volumes. D) Geometric mean calcification surface / volume ratio. E) Geometric mean sphericity. F) Geometric mean closed porosity. For closed porosity, only samples that had particles large enough to have closed porosity were included in the unidentified group.

**Table II.** Quantitative parameters for BCP, CPP, and their difference presented as geometric means.

| Parameter | BCP<br>(geometric mean) | CPP<br>(geometric mean) | Ratio between CPP vs BCP<br>(95% CI) |
| --- | --- | --- | --- |
| Total calcification volume (mm<sup>3</sup>) | 0.075 | 1.014 | 14 ( 3 - 73) |
| Total number of particles | 1178 | 8716 | 7 ( 3 - 17) |
| Average particle volume (μm<sup>3</sup>) | 9419 | 37650 | 7 ( 2 - 8) |
| Surface to volume ratio | 0.845 | 0.983 | 1.16 ( 1.08 - 1.25) |
| Sphericity | 0.698 | 0.624 | 0.89 ( 0.83 - 0.95) |
| Closed porosity (%) | 9.87 | 3.76 | 0.38 ( 0.23 - 0.63) |

## 4. Discussion

This study investigated calcification patterns in human posterior horn menisci *ex vivo* using high-resolution 3D µCT imaging. Morphological and volumetric differences were identified between BCP and CPP deposits. BCP calcifications were typically located on meniscal surfaces and near tears, while CPP deposits were embedded within the tissue, aligned with circumferential collagen fibers. Quantitative analysis revealed that BCP deposits had lower total volume, fewer and smaller particles, and lower surface-to-volume ratios compared to CPP, whereas BCP showed higher sphericity and closed porosity. These findings demonstrate that µCT imaging enables reliable differentiation between BCP and CPP calcifications in human meniscus.

Our findings suggest that BCP accumulation and morphology may play a direct role in meniscal degradation in OA. Qualitatively, BCP crystals were primarily located on the surface, periphery, and within complex meniscal tears. Given the calcifying potential of OA meniscal cells and the absence of intact surrounding tissue, some particles may have migrated through joint cavity from other joint structures, such as articular cartilage, where calcified crystals are also commonly found^18,19^. BCPs appeared in punctate patterns but formed sharp-edged clusters when aggregated. These clusters may influence biomechanical properties of the tissue contribute to mechanical degradation during joint movement, embedding into fibrillated meniscal tissue and advancing degeneration. Previous studies have linked BCPs to destructive processes in conditions like Milwaukee shoulder syndrome and intraosseous migration, and intra-articular mineralization has been associated with knee pain^20,21^.

In our samples, CPP calcifications were predominantly located within the meniscus, forming large rod-like structures aligned with the circumferential collagen bundles. In some cases, smaller, discontinuous clusters were observed in similar orientations, potentially representing precursors to the rod-like form. The smooth, elongated morphology of these calcifications suggests they occupy fluid channels between collagen fibers, consistent with known pathways where fluid flows during biomechanical loading^22,23^. Furthermore, CPPs were found on meniscal surfaces and tears as amorphous, randomly oriented deposits, distinct from the organized rods inside the meniscus. Surface-to-volume analysis revealed CPPs have more complex surfaces, while BCP particles were more spherical. Notably, BCPs exhibited a porous structure that was absent in CPPs, as confirmed by both qualitative and quantitative porosity assessments.

CPP calcifications in the meniscus may be associated with vascular structures, as seen in 3D µCT images showing complex, curved calcifications oriented nonlinearly toward tibial or femoral surfaces. Hollow, vessel-like calcified structures suggest vascular involvement. This aligns with prior findings linking arterial calcifications and atherosclerosis to OA^24^, and calcification clusters have been reported to form near the largest vascular supply in the outer region of porcine meniscus^15^. The shape of these calcified hollow structures resembles meniscal vascularity that has been studied previously with µCT^15,25^. The shape of these structures resembles previously characterized meniscal vasculature but was observed only in CPP samples, while arterial calcifications have been reported to typically contain hydroxyapatite^26^. The absence of small particles inside these structures suggests that CPP likely forms locally from excess calcium ions that combine with pyrophosphate to form calcium pyrophosphate dihydrate crystals rather than being transported via the vasculature^27,28^.

The closed porosity of BCP deposits consisted of organized porous structures inside the calcifications, which was not observed in CPP deposits to similar extent. Accurate differentiation of BCP and CPP calcifications in the soft tissues has been challenging with current imaging methods. Previous characterizing methods, like Raman and SEM-ED, require destructive sample processing, while current imaging methods, like MECT may not have sufficient resolution and accuracy^29–31^. The organized network of porous structures in BCP calcifications could therefore be used as an imaging biomarker for characterization of BCP from CPP calcifications with high-resolution imaging modalities like µCT and MECT combined with PCDs in future studies.

Based on the 3D µCT images and videos, we identified important differences between the BCP and CPP calcification patterns. As previously stated, BCPs were smaller, punctate, and found in the peripheral area, surface, and tears, spread along evenly with few larger clusters. While some amorphous CPP were on the surfaces of samples, the largest volumes of CPP were located inside the meniscus, along the circumferential collagen fibers, forming long rod-like formations. Similarly, a previous study has reported that in 2D, CPP formed round colonies that disrupt the collagen fiber organization^32^. Furthermore, BCPs were found both qualitatively and quantitatively in smaller volumes, fewer in number, and smaller in average particle size compared to CPP. Previously these aggregates of BCP and CPP have been suggested to be approximately between 1 to 20µm in diameter^33^. For comparison, our volumetric results for BCP and CPP were 9419µm^3^ and 37650µm^3^, which can produce approximated sphere diameters of 26µm and 42µm, respectively, which are higher than previously reported diameters from histological sections. The discrepancy between the previous studies and our study can be from our results including 3D information from larger samples.

Meniscal calcification may be linked to danger-associated molecular patterns (DAMPs) in the knee joint, as microcrystals like BCP can trigger pro-inflammatory responses and promote chondrocyte hypertrophy^34,35^. We saw positive Safranin O staining which marks an increase in proteoglycan content and hypertrophic cells in the vicinity of BCP calcifications. Furthermore, BCP has been shown to induce hypertrophic differentiation^36–38^, a key step in endochondral ossification, where cartilage transitions to bone through chondrocyte proliferation, hypertrophy, and apoptosis^39,40^. Apoptotic chondrocytes are associated with BCP crystals and may contribute to OA via calcifying apoptotic bodies^41,42^. Therefore, a similar fate of meniscal cells undergoing calcification may happen in the meniscus to hypertrophic chondrocyte-like cells that are near BCP deposits. It has been suggested that similar molecular mechanisms that control regular calcification in skeletal tissues are similar to calcification mechanisms in soft tissues^43^. Our µCT analysis revealed that BCP calcifications form porous, lacunae-like structures resembling osteocyte organization in subchondral bone and calcified cartilage. This pattern was consistently observed in 3D across all BCP samples but absent in CPP calcifications, indicating distinct calcification mechanisms. The unique architecture of BCP may be attributed to its high hydroxyapatite content, which plays a key role in bone regeneration^44^. These findings suggest that BCP accumulation in the meniscus mimics native bone tissue formation, supporting the hypothesis that soft tissue calcification may share molecular pathways with physiological skeletal mineralization.

We found only a few cases which had both calcification types in the same menisci, and in those cases, they were observed in different areas. In those samples, BCPs covered a relatively low volume compared to the whole calcification volume. Likewise, there has been incidents where both crystal types have been observed in articular cartilage^45^. CPP crystal formation is closely associated with the regulation of extracellular inorganic pyrophosphate, which also inhibits BCP crystal growth^46^. Dysregulation of pyrophosphate can be related to aging or genetic mutations, which leads to increased pyrophosphate crystal deposition^47^. CPP calcifications have been associated with senescent chondrocytes^36^. The cells around solid, rod-like CPP calcifications were elliptic-shaped, while cells around amorphous CPP calcifications seemed more hypertrophic. As the surface of solid CPP is smooth and arranged along the circumferential collagen fibers, this could minimize the disrupting effect the calcification cluster has on the meniscus function and account for the asymptomatic CPPD disease. Previous studies have shown increase in type X collagen near calcified fibrocartilage of meniscus, which acts as a facilitator and regulator for matrix mineralization^48,49^. This could suggest that after initial inflammation and formation of the calcification inside the meniscus, the smooth calcification particle does not severely disrupt the surrounding meniscal ECM.

Calcifications were present in all OA menisci, with basic calcium phosphate (BCP) being the predominant type and correlated with Pauli histological degeneration score. ^9^. Consistently, higher calcification grades were associated with increased Pauli scores^5^. Notably, samples with CPP deposits exhibited the highest average calcification grades. A minimum grade of 3 was required for calcifications to be detectable in histological sections or via Raman spectroscopy, with grade 3 deposits approximating a total volume of 0.005 mm³ or 0.01% of total meniscus volume. Although only 8 of 74 samples lacked visible calcifications on qualitative 3D µCT assessment, all samples contained minute particles in volumetric analysis, suggesting early microcrystal formation. These findings in OA-free menisci add to previous reports of widespread calcifications in OA meniscus^50^, articular cartilage^18,45,51^, and synovial fluid^52^. Given the average age of 71 and mild to moderate degeneration in our cohort, our results support the notion that minor calcifications are common with aging and probably BCPs^5,51^. Thus, future studies should aim at quantifying what volume of calcifications could be indicative of OA disease. Potential OA therapies could consider calcification inhibition as a potential disease-modifying strategy.

A limitation of the study was that only a portion of the meniscus posterior horn was imaged, which enabled optimal resolution due to µCT’s geometrical magnification constraints. Despite the limited size, the imaged volume was substantially larger than in conventional 2D histology.

This study lays the groundwork on how the calcification types can be differentiated by their calcification patterns and quantitative differences in 3D using µCT. Differences in total volume, number of particles, individual particle volume, calcification surface to volume ratio, sphericity, and especially in porosity were observed with µCT imaging in the calcification patterns between meniscal BCP and CPP calcifications. The porosity marker could help in differentiating BCP and CPP calcification with sensitive enough 3D imaging modalities in the future to lessen the requirement of spectroscopy or other highly specialized modalities. This quantitative comparison of meniscal calcification patterns may provide a better understanding regarding the role of calcifications in OA.

## Acknowledgements

We would like to thank the MENIX clinical staff at Trelleborg Hospital, the Tissue Donor Bank at Skåne University Hospital, and the Department of Forensic Medicine in Lund for their collaboration that enabled sample collection. We would also like to acknowledge Piia Mäkelä, Laboratory Technician, for the preparatory work with the histological samples.

## Author contributions

Conception and design: VPK, IH, SS, ME, AT. Provision of study materials and tissue preparation: ME, VH, PÖ, JT, VPK, KE, AS, SS. Micro-computed tomography imaging and analysis: VPK, EN. Raman spectroscopy and analysis: BS. Histological analysis: VPK, IH, NK, SDG. Statistical analysis: AT, VPK. Interpretation of results: All coauthors. Drafting of the article: VPK. Critical revision of the article for important intellectual content: All coauthors, Final approval of the article: All coauthors.

## References

1. Pauli C, Grogan SP, Patil S, et al. Macroscopic And Histopathologic Analysis Of Human Knee Menisci In Aging And Osteoarthritis. Osteoarthritis Cartilage. 2011;19(9):1132–1141. doi:10.1016/j.joca.2011.05.008

2. Sun Y, Mauerhan DR, Honeycutt PR, et al. Calcium deposition in osteoarthritic meniscus and meniscal cell culture. Arthritis Res Ther. 2010;12(2):R56. doi:10.1186/AR2968

3. MacMullan PA, McCarthy GM. The meniscus, calcification and osteoarthritis: A pathologic team. Arthritis Res Ther. 2010;12(3):1–2. doi:10.1186/AR2993/METRICS

4. Sun Y, Mauerhan DR. Meniscal calcification, pathogenesis and implications. Curr Opin Rheumatol. 2012;24(2):152–157. doi:10.1097/BOR.0B013E32834E90C1

5. Hellberg I, Karjalainen VP, Finnilä MAJ, et al. 3D analysis and grading of calcifications from ex vivo human meniscus. Osteoarthritis Cartilage. 2023;31(4):482–492. doi:10.1016/J.JOCA.2022.10.016

6. Nasi S, So A, Combes C, Daudon M, Busso N. Interleukin-6 and chondrocyte mineralisation act in tandem to promote experimental osteoarthritis. Ann Rheum Dis. 2016;75(7):1372–1379. doi:10.1136/ANNRHEUMDIS-2015-207487

7. Cheung HS, Sallis JD, Demadis KD, Wierzbicki A. Phosphocitrate blocks calcification-induced articular joint degeneration in a guinea pig model. Arthritis Rheum. 2006;54(8):2452–2461. doi:10.1002/ART.22017

8. Bernabei I, So A, Busso N, Nasi S. Cartilage calcification in osteoarthritis: mechanisms and clinical relevance. Nature Reviews Rheumatology 2022 19:1. 2022;19(1):10–27. doi:10.1038/s41584-022-00875-4

9. Shakya BR, Karjalainen VP, Hellberg I, et al. Prevalence and classification of meniscal calcifications in the human knee. Osteoarthritis Cartilage. 2024;32(11):1443–1451. doi:10.1016/J.JOCA.2024.07.013

10. Ea HK, Chobaz V, Nguyen C, et al. Pathogenic Role of Basic Calcium Phosphate Crystals in Destructive Arthropathies. PLoS One. 2013;8(2):e57352. doi:10.1371/JOURNAL.PONE.0057352

11. Fuerst M, Bertrand J, Lammers L, et al. Calcification of articular cartilage in human osteoarthritis. Arthritis Rheum. 2009;60(9):2694–2703. doi:10.1002/ART.24774

12. Nevanranta E, Karjalainen VP, Brix M, et al. Characterizing meniscal calcifications with photon counting-based dual-energy computed tomography. bioRxiv. Published online February 18, 2025:2025.02.14.638040. doi:10.1101/2025.02.14.638040

13. Stamp LK, Anderson NG, Becce F, et al. Clinical Utility of Multi-Energy Spectral Photon-Counting Computed Tomography in Crystal Arthritis. Arthritis and Rheumatology. 2019;71(7):1158–1162. doi:10.1002/ART.40848/ABSTRACT

14. Karjalainen VP, Kestilä I, Finnilä MA, et al. Quantitative three-dimensional collagen orientation analysis of human meniscus posterior horn in health and osteoarthritis using micro-computed tomography. Osteoarthritis Cartilage. 2021;29(5). doi:10.1016/j.joca.2021.01.009

15. Karjalainen VP, Herrera Millar VR, Modina | Silvia, et al. Age and anatomical region-related differences in vascularization of the porcine meniscus using microcomputed tomography imaging. Journal of Orthopaedic Research®. Published online April 29, 2024. doi:10.1002/JOR.25862

16. Kestilä I, Folkesson E, Finnilä MA, et al. Three-dimensional microstructure of human meniscus posterior horn in health and osteoarthritis. Osteoarthritis Cartilage. 2019;27(12):1790–1799. doi:10.1016/j.joca.2019.07.003

17. Sergio M, Karjalainen VP, Das Gupta S, et al. Prolonged excessive weight induces spontaneous meniscal degeneration in sows: A preclinical model for obesity-related knee OA. Annals of Anatomy - Anatomischer Anzeiger. 2025;260:152681. doi:10.1016/J.AANAT.2025.152681

18. Stücker S, Koßlowski F, Buchholz A, Lohmann CH, Bertrand J. High frequency of BCP, but less CPP crystal-mediated calcification in cartilage and synovial membrane of osteoarthritis patients. Osteoarthritis Cartilage. Published online May 11, 2024. doi:10.1016/J.JOCA.2024.04.019

19. Kiraly AJ, Roberts A, Cox M, Mauerhan D, Hanley E, Sun Y. Comparison of Meniscal Cell-Mediated and Chondrocyte-Mediated Calcification. Open Orthop J. 2017;11(1):225. doi:10.2174/1874325001711010225

20. Halverson PB, Cheung HS, Mccarty DJ, Garancis J, Mandel N. “Milwaukee shoulder”—association of microspheroids containing hydroxyapatite crystals, active collagenase, and neutral protease with rotator cuff defects. ii. synovial fluid studies. Arthritis Rheum. 1981;24(3):474–483. doi:10.1002/ART.1780240304

21. Collinot JA, Pascart T, Budzik JF, Hügle T, Hussenot M, Becce F. Non-invasive characterization of intra-articular mineralization using dual-energy computed tomography. Rheumatology. 2020;59(12):3997–3998. doi:10.1093/RHEUMATOLOGY/KEAA231

22. Agustoni G, Maritz J, Kennedy J, Bonomo FP, Bordas SPA, Barrera O. High Resolution Micro-Computed Tomography Reveals a Network of Collagen Channels in the Body Region of the Knee Meniscus. Ann Biomed Eng. 2021;49(9):2273–2281. doi:10.1007/S10439-021-02763-6/FIGURES/4

23. Vetri V, Dragnevski K, Tkaczyk M, et al. Advanced microscopy analysis of the micro-nanoscale architecture of human menisci. Sci Rep. 2019;9(1). doi:10.1038/s41598-019-55243-2

24. Macêdo MB, Santos VMOS, Pereira RMR, Fuller R. Association between osteoarthritis and atherosclerosis: A systematic review and meta-analysis. Exp Gerontol. 2022;161. doi:10.1016/J.EXGER.2022.111734

25. Orellana F, Grassi A, Hlushchuk R, et al. Revealing the complexity of meniscus microvasculature through 3D visualization and analysis. Scientific Reports 2024 14:1. 2024;14(1):1–14. doi:10.1038/s41598-024-61497-2

26. Villa-Bellosta R. Synthesis of extracellular pyrophosphate increases in vascular smooth muscle cells during phosphate-induced calcification. Arterioscler Thromb Vasc Biol. 2018;38(9):2137–2147. doi:10.1161/ATVBAHA.118.311444/SUPPL_FILE/ATVB_ATVB-2018-311444D_SUPP2.PDF

27. Kosik-Bogacka DI, Lanocha-Arendarczyk N, Kot K, et al. Calcium, magnesium, zinc and lead concentrations in the structures forming knee joint in patients with osteoarthritis. Journal of Trace Elements in Medicine and Biology. 2018;50:409–414. doi:10.1016/J.JTEMB.2018.08.007

28. Habata T, Ohgushi H, Takakura Y, et al. Relationship between meniscal degeneration and element contents. Biol Trace Elem Res. 2001;79(3):247–256. doi:10.1385/BTER:79:3:247/METRICS

29. Bernabei I, Sayous Y, Raja AY, et al. Multi-energy photon-counting computed tomography versus other clinical imaging techniques for the identification of articular calcium crystal deposition. Rheumatology. 2021;60(5):2483–2485. doi:10.1093/RHEUMATOLOGY/KEAB125

30. Budzik JF, Marzin C, Legrand J, Norberciak L, Becce F, Pascart T. Can Dual-Energy Computed Tomography Be Used to Identify Early Calcium Crystal Deposition in the Knees of Patients With Calcium Pyrophosphate Deposition? Arthritis and Rheumatology. 2021;73(4):687–692. doi:10.1002/ART.41569/ABSTRACT

31. Jarraya M, Bitoun O, Wu D, et al. Dual energy computed tomography cannot effectively differentiate between calcium pyrophosphate and basic calcium phosphate diseases in the clinical setting. Osteoarthr Cartil Open. 2024;6(1):100436. doi:10.1016/J.OCARTO.2024.100436

32. Katsamenis OL, Karoutsos V, Kontostanos K, Panagiotopoulos EC, Papadaki H, Bouropoulos N. Microstructural characterization of CPPD and hydroxyapatite crystal depositions on human menisci. Crystal Research and Technology. 2012;47(11):1201–1209. doi:10.1002/CRAT.201200346

33. Schumacher HR. Crystal-induced arthritis: An overview. Am J Med. 1996;100(2):46S–52S. doi:10.1016/S0002-9343(97)89546-0

34. McAllister MJ, Chemaly M, Eakin AJ, Gibson DS, McGilligan VE. NLRP3 as a potentially novel biomarker for the management of osteoarthritis. Osteoarthritis Cartilage. 2018;26(5):612–619. doi:10.1016/J.JOCA.2018.02.901

35. Busso N, So A. Microcrystals as DAMPs and their role in joint inflammation. Rheumatology. 2012;51(7):1154–1160. doi:10.1093/RHEUMATOLOGY/KER524

36. Meyer F, Dittmann A, Kornak U, et al. Chondrocytes From Osteoarthritic and Chondrocalcinosis Cartilage Represent Different Phenotypes. Front Cell Dev Biol. 2021;9:622287. doi:10.3389/FCELL.2021.622287/BIBTEX

37. Dreier R. Hypertrophic differentiation of chondrocytes in osteoarthritis: The developmental aspect of degenerative joint disorders. Arthritis Res Ther. 2010;12(5):1–11. doi:10.1186/AR3117/TABLES/1

38. Bertrand J, Kräft T, Gronau T, et al. BCP crystals promote chondrocyte hypertrophic differentiation in OA cartilage by sequestering Wnt3a. Ann Rheum Dis. 2020;79(7):975–984. doi:10.1136/ANNRHEUMDIS-2019-216648

39. Mackie EJ, Ahmed YA, Tatarczuch L, Chen KS, Mirams M. Endochondral ossification: How cartilage is converted into bone in the developing skeleton. Int J Biochem Cell Biol. 2008;40(1):46–62. doi:10.1016/J.BIOCEL.2007.06.009

40. Aghajanian P, Mohan S. The art of building bone: emerging role of chondrocyte-to-osteoblast transdifferentiation in endochondral ossification. Bone Research 2018 6:1. 2018;6(1):1–9. doi:10.1038/s41413-018-0021-z

41. Kourí JB, Aguilera JM, Reyes J, Lozoya KA, González S. Apoptotic chondrocytes from osteoarthrotic human articular cartilage and abnormal calcification of subchondral bone. J Rheumatol. 2000;27(4):1005–1019. Accessed June 11, 2025. https://europepmc.org/article/med/10782830

42. Chen WH, Lo WC, Hsu WC, et al. Synergistic anabolic actions of hyaluronic acid and platelet-rich plasma on cartilage regeneration in osteoarthritis therapy. Biomaterials. 2014;35(36):9599–9607. doi:10.1016/J.BIOMATERIALS.2014.07.058

43. Yan J fei, Qin W pin, Xiao B cheng, et al. Pathological calcification in osteoarthritis: an outcome or a disease initiator? Biological Reviews. 2020;95(4):960–985. doi:10.1111/BRV.12595

44. Kattimani VS, Kondaka S, Lingamaneni KP. Hydroxyapatite–-Past, Present, and Future in Bone Regeneration. Bone Tissue Regen Insights. 2016;7:BTRI.S36138. doi:10.4137/BTRI.S36138

45. Nguyen C, Bazin D, Daudon M, et al. Revisiting spatial distribution and biochemical composition of calcium-containing crystals in human osteoarthritic articular cartilage. Arthritis Res Ther. 2013;15(5):1–12. doi:10.1186/AR4283/TABLES/3

46. Johnson K, Terkeltaub R. Inorganic pyrophosphate (PPI) in pathologic calcification of articular cartilage. Front Biosci. 2005;10:988–997. doi:10.2741/1593

47. Zaka R, Williams CJ. Genetics of chondrocalcinosis. Osteoarthritis Cartilage. 2005;13(9):745–750. doi:10.1016/J.JOCA.2005.04.006

48. Shen G, Shen G. The role of type X collagen in facilitating and regulating endochondral ossification of articular cartilage. Orthod Craniofac Res. 2005;8(1):11–17. doi:10.1111/J.1601-6343.2004.00308.X

49. Iiro Eerola, Heli Salminen, Pirkko Lammi, et al. Type X collagen, a natural component of mouse articular cartilage: Association with growth, aging, and osteoarthritis. 1998;41(7):1287–1295. doi:10.1002/1529-0131(199807)41:7

50. Mitsuyama H, Healey RM, Terkeltaub RA, Coutts RD, Amiel D. Calcification of human articular knee cartilage is primarily an effect of aging rather than osteoarthritis. Osteoarthritis Cartilage. 2007;15(5):559–565. doi:10.1016/J.JOCA.2006.10.017

51. Derfus BA, Kurian JB, Butler JJ, et al. The high prevalence of pathologic calcium crystals in pre-operative knees. J Rheumatol. 2002;29(3).

