## Supplementary materials for "Characterization of calcifications in posterior horn of human meniscus using micro-computed tomography"

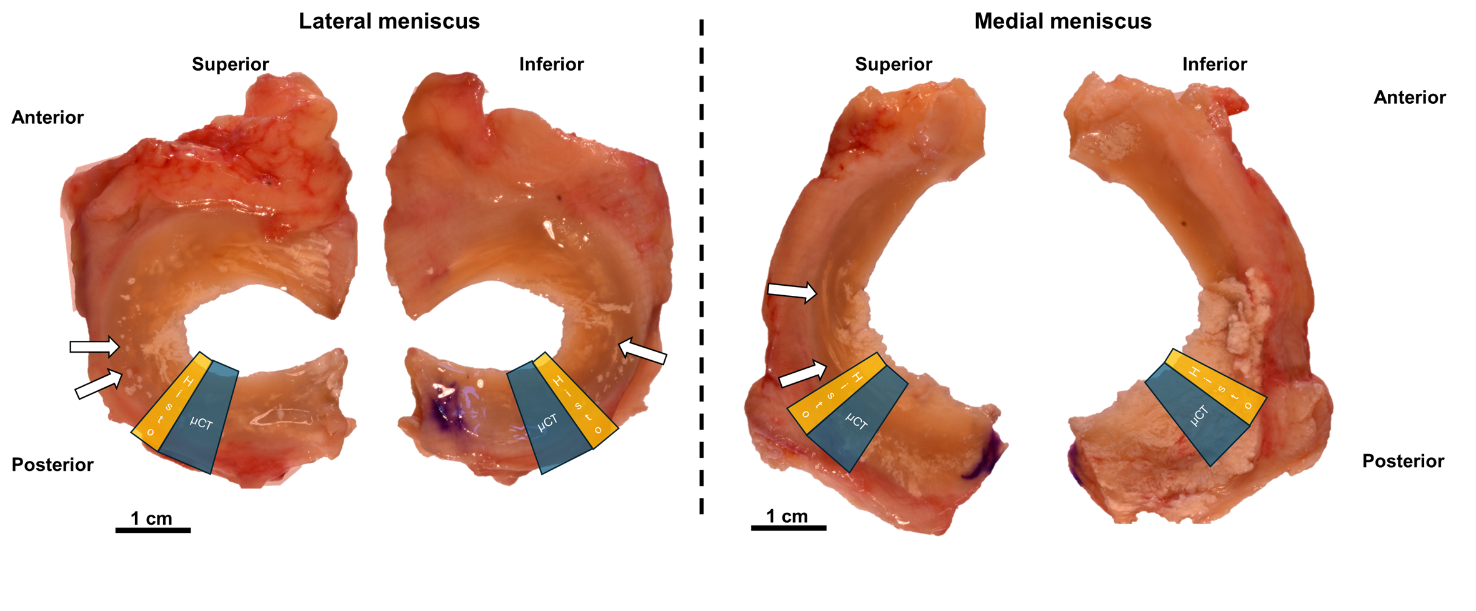

Supplementary Figure S1. An example of deceased donor’s lateral and medial meniscus. Rod-like calcium pyrophosphate (CPP) calcifications can be seen inside the tissue (arrows), following the circumferential shape of meniscus. Location of the micro-computed tomography (µCT) piece is shown in blue. Location of cut sections for histological analysis and Raman spectroscopy is shown in yellow.

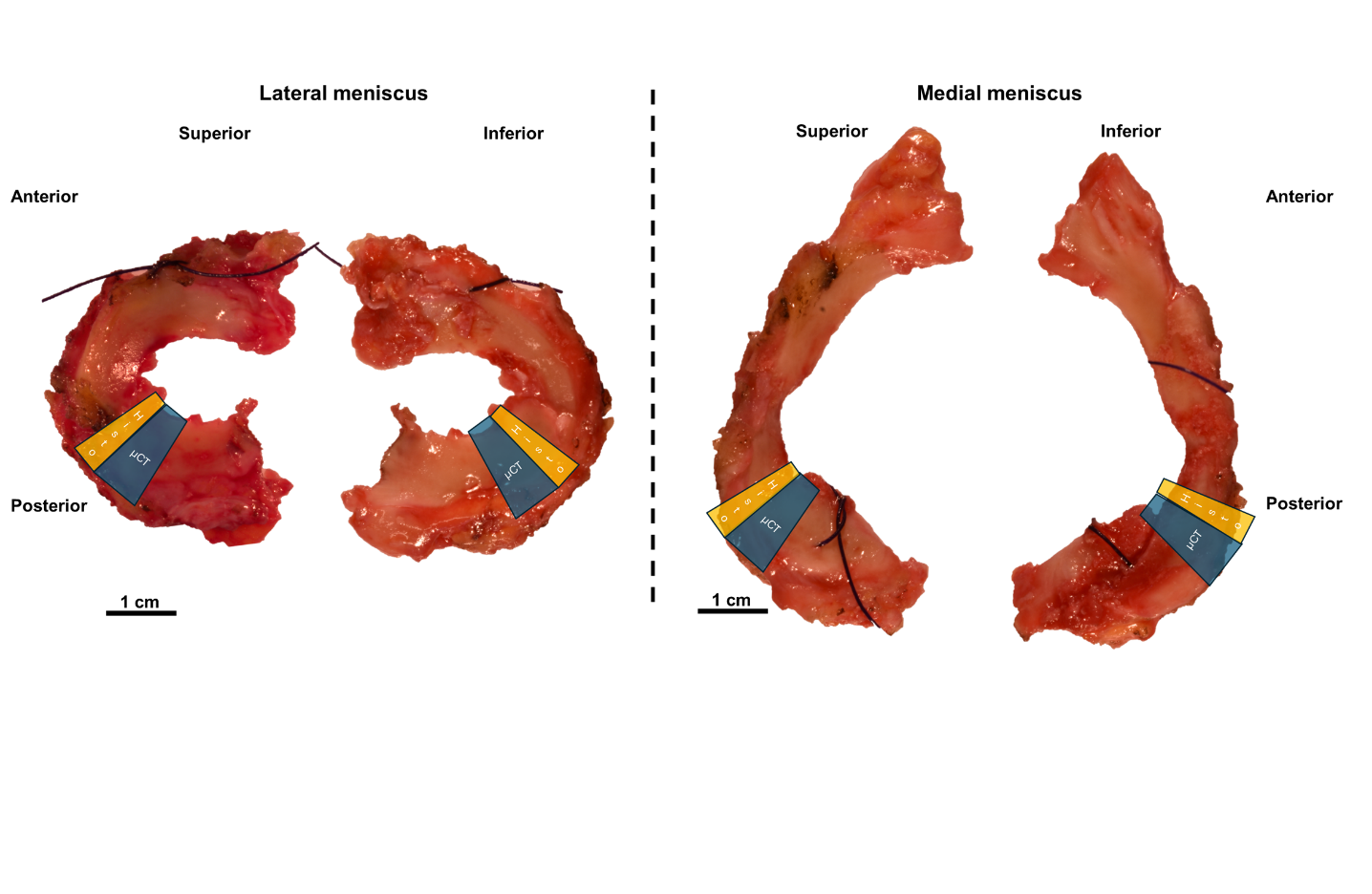

Supplementary Figure S2. An example of a lateral and medial meniscus from a total knee replacement patient with medial compartment osteoarthritis. No visible calcifications are seen. Location of the micro-computed tomography piece is shown in blue. Location of histological sections and Raman spectroscopy is shown in yellow.

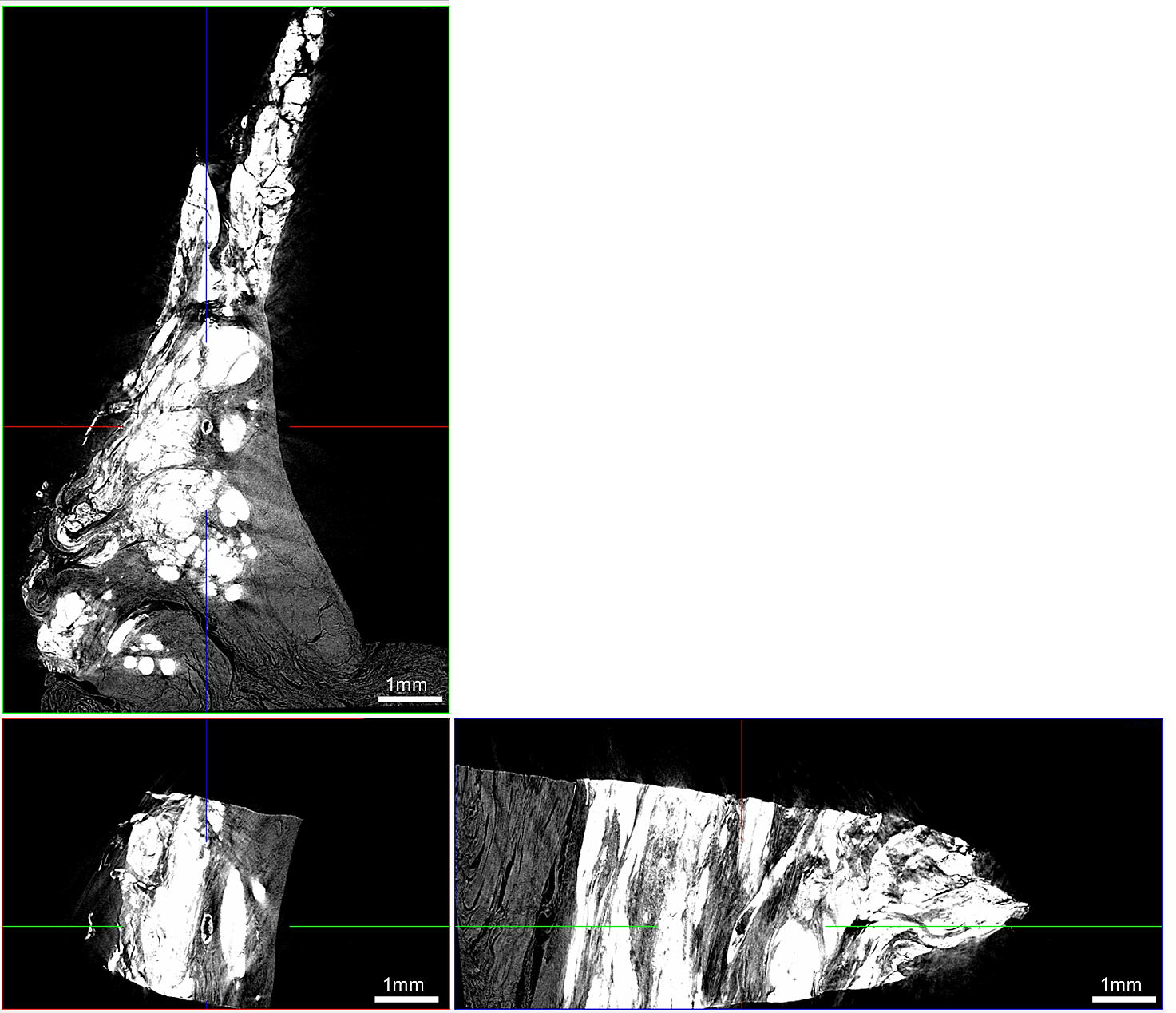

Supplementary Figure S3. An example cross-section from three directions of a hollow rod-shaped calcification in a sample with identified CPP calcifications.

Supplementary Material IV

Description of calcifications, cells, and proteoglycan content,

An example meniscus sample with both BCP (blue arrow) and CPP (red arrow) calcifications identified with Raman spectroscopy is shown in supplementary figure S4. The BCP calcifications in general were found attached to the surface, fibrillations, or periphery of the meniscus. Figure S4B shows distinct shape and density differences between BCP and CPP aggregates, with BCP appearing denser and CPP accumulating in rod-like shapes, covering the whole sample from the anterior to the posterior side, while BCP forms clusters with sharp edges. In Figure S4C, adjacent histology sections shows BCP (blue arrow) only as a small cluster in the periphery of the meniscus outer region. CPP (red arrow and all neighboring calcifications) are seen inside the meniscus and near teared area. In Figure S4E, the left arrow shows amorphous CPP calcification, while the right arrow shows solid, circumferential rod-like CPP. In Figure S4G, BCP is located in the periphery of the outer region of meniscus, with high cell count in the joint capsule with vascular supply and only a few cells in the outer region. In Figure S4H, the cells around solid CPP are ellipsoid-shaped and oriented along the calcification surface, similar to chondrocytes in articular cartilage. Importantly: there are no distinct cells or lacunae structures or staining visible inside the BCP or CPP calcifications in histological sections (Supplementary figure S4 H&K). Safranin O staining intensity is increased in the whole sample (Supplementary figure S4I). However, the areas near solid CPPs are more pronounced in staining, compared to more amorphous CPP areas (Supplementary figure S4 J&K).
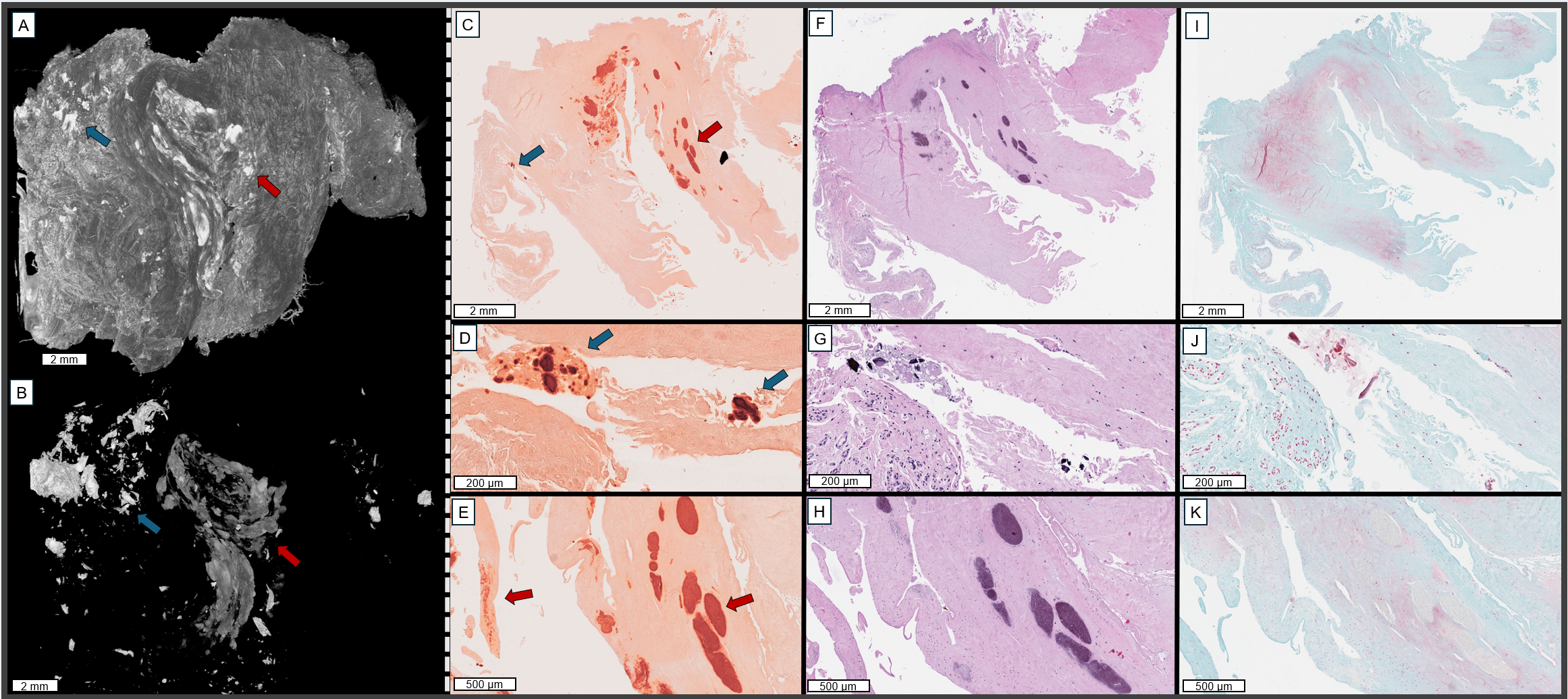
Supplementary figure S4. A) 3D µCT image of OA meniscus with both BCP (blue arrow) and CPP (red arrow) calcifications, as identified with Raman spectroscopy measurements of an adjacent tissue section. B) 3D µCT image of the calcifications without soft tissue. C-E) Alizarin Red -stained histological section adjacent to the µCT piece. C) BCP highlighted with blue arrow and CPP with red arrow. D) Magnified image of BCP from 3C in slightly different orientation. E) Magnified image from 3C of CPP with amorphous CPP on the left and solid, ellipsoidal CPP on the right. F, G, H) Adjacent hematoxylin and eosin -stained sections show cell nuclei near calcifications. I, J, K) Adjacent Safranin O - Fast Green -stained sections show increased proteoglycan staining in the whole tissue, while solid CPP areas have more Safranin O staining around them compared to the surroundings of amorphous CPP areas.

Supplementary figure S5 A&B shows that the particles are located in the complex surface fibrillations and tears of the meniscal surface of a meniscus sample from an individual with OA inhabiting only BCP calcification. From histological sections in Figure S4C-E, the complex 3D tears are not visible and the BCP particles are seemingly inside the sample. In Figure S4F-H, the cells seem hypertrophic, and only a few normal, healthy cells are seen around the BCP aggregates. Safranin O staining in Figures S4I-K reveals increased staining in the whole sample. Similar to the sample in Figure S4, the BCP calcifications do not show distinct porosity inside their structure in histological sections.

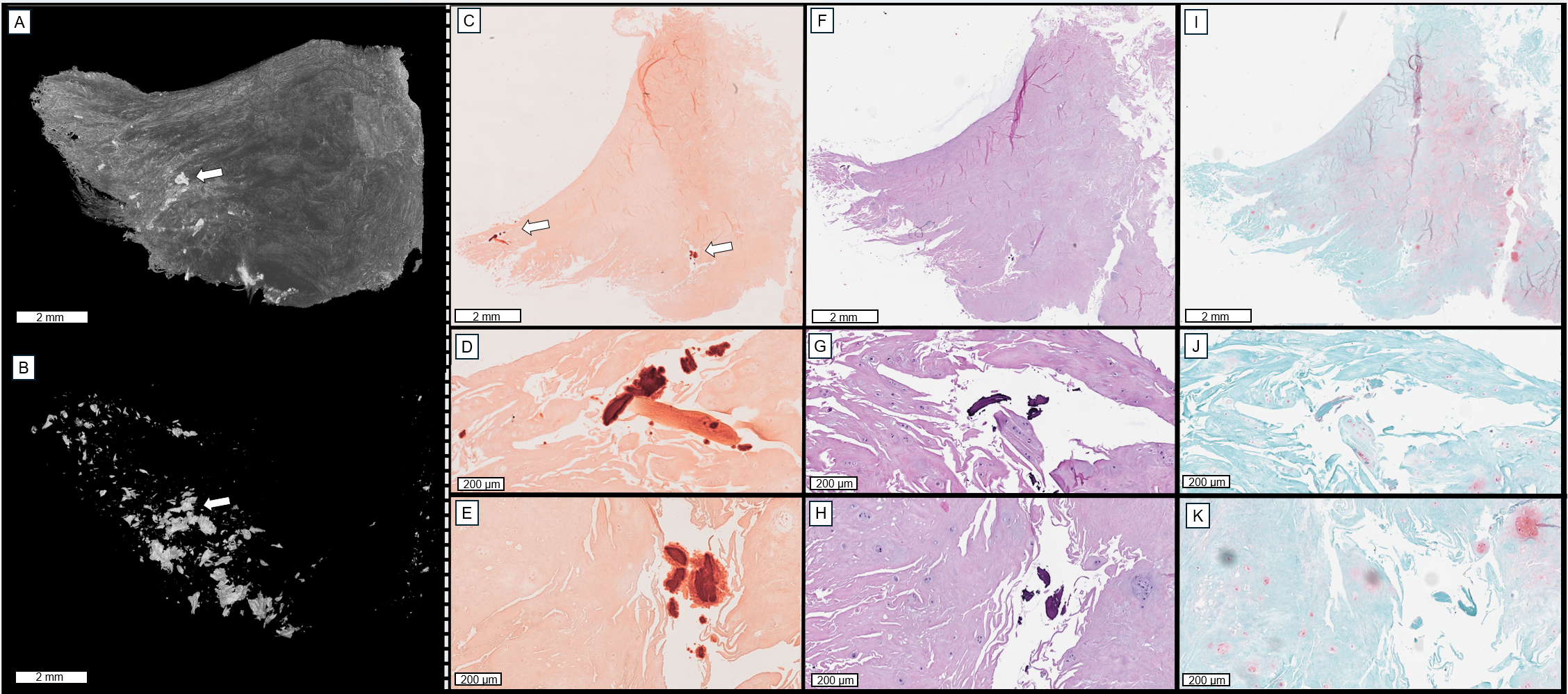

Supplementary figure S5. A) 3D µCT image of a meniscus sample from an individual with OA inhabiting only BCP calcifications. B) The calcifications are inside the fibrillations and tears near the meniscus surface, and a few punctate calcifications are seen on the outer region. C, D, E) Alizarin red staining shows two distinct calcification clusters that are seemingly inside the meniscus. F, G, H) Hematoxylin and eosin staining shows few hypertrophic cells and loss of healthy cells around the tear, where BCP calcifications are located. I, J, K) Safranin O - Fast Green staining shows an increase in proteoglycan content and oedema-like features in the sample.

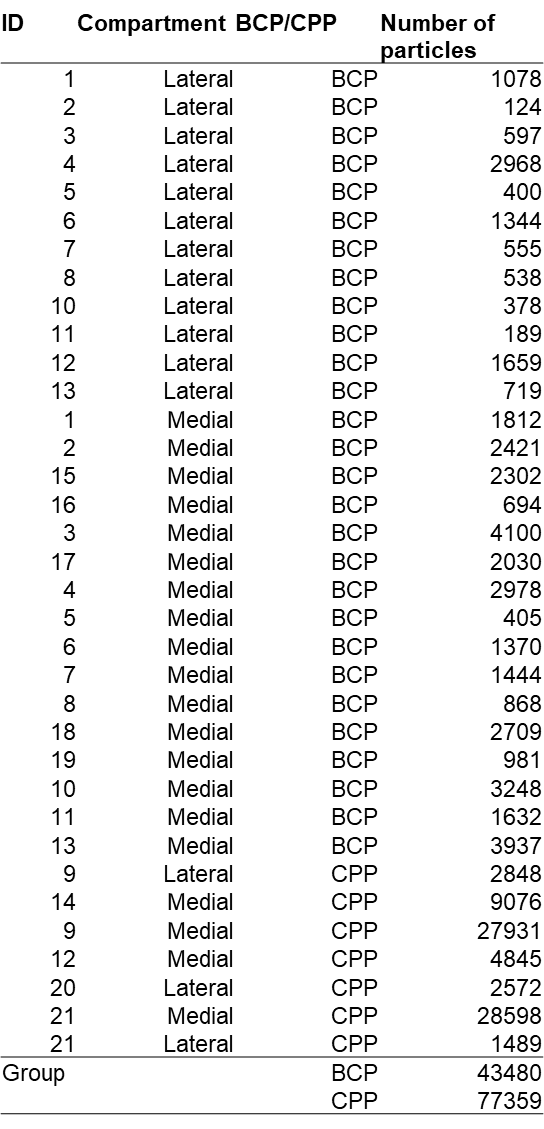

Supplementary Figure S6. Total number of analyzed particles for each meniscus.
